# Influence of age and cognitive performance on resting-state functional connectivity of dopaminergic and noradrenergic centres

**DOI:** 10.1101/2022.02.27.482151

**Authors:** Michal Rafal Zareba, Wiktoria Furman, Marek Binder

## Abstract

The process of ageing is associated with structural and functional changes in the brain, with a decline in cognitive functions observed as its inevitable symptom, even in the absence of neurodegenerative changes. A body of literature suggests dopamine and noradrenaline as prominent candidate molecules to mediate these effects, however, the knowledge regarding the underlying mechanisms is scarce. To fill this gap, we compared local and distant resting-state functional connectivity patterns of ventral tegmental area, substantia nigra pars compacta and locus coeruleus in healthy young (20-35 years; N = 37) and older adults (55-80 years; N = 27). Additionally, we sought long-range functional connectivity patterns of these structures associated with performance in tasks probing executive, attentional and reward functioning, as well as compared the functional coupling of the left and right substantia nigra. The results showed that each substantia nigra had stronger coupling with ipsilateral cortical and subcortical areas along with contralateral cerebellum, and that strength of functional connections of this structure with angular gyrus and lateral orbitofrontal cortex predicted the visuomotor search abilities. In turn, ageing was associated with increased local synchronisation in the ventral tegmental area, and differences in functional connectivity of the dopaminergic midbrain with numerous cortical and subcortical structures. Locus coeruleus functional coupling showed no differences between the groups and was not associated with any of the behavioural functions. To the best of our knowledge, this work is the first one to report the age-related effects on midbrain local synchronisation and its connectivity with key recipients of dopaminergic innervation, such as striatum, anterior cingulate and insula.

## 1. Introduction

The recent decades have seen the developed countries undergo dramatic changes in their demographic structure. An increase in the quality of life and high availability of medical services has resulted in the lengthening of life expectancy. Ageing is known to be associated with the increased risk of the development of neurodegenerative diseases. Even in the absence of pathological alterations, substantial changes in the structure and functioning of the human brain occur (Tsvetanov et al., 2021), and a prominent decline in cognitive functions such as attention, executive functioning, working memory, conceptual reasoning and visuospatial processing is observed (Hedden and Gabrieli, 2004; Harada et al., 2013). Interestingly, certain other processes, such as reward responsiveness, remain intact in the elderly individuals (Samanez-Larkin and Knutson, 2015). The ability to discriminate between physiological changes associated with ageing and pathological alterations is essential for the early detection of signs of neurodegenerative diseases and the establishment of particular risk factors. Therefore, this topic has been widely studied both in human and animal models.

Resting-state magnetic functional imaging (rs-fMRI) constitutes a noninvasive method that enables in vivo investigation of changes in the functioning of the human brain. Resting-state functional connectivity is founded on the phenomenon of temporal correlations of activation signals between anatomically separated brain regions that reflect both direct and indirect structural connectivity (Vincent et al., 2007). The functional connectivity approach provides information regarding the strength of signal correlation between distanced brain structures and their networks, while regional homogeneity (ReHo) offers an insight into local synchronisation and functional specificity (Zang et al., 2004). There has been a notable increase in the number of rs-fMRI studies related to ageing in the last two decades. Results from different studies referring to age-related changes are not congruent, for both decreases and increases in resting-state functional networks were observed. The most consistent findings across different studies are a decline in functional connectivity within networks and increased functional connectivity between networks (Jockwitz and Caspers, 2021). However, the majority of studies dedicated to age-related changes in connectivity focused on the main known resting-state networks, such as the default mode network (DMN) and attentional networks. Many brain centres related to functions affected by ageing do not belong to these main functional networks, including catecholaminergic nuclei in the midbrain and brainstem that are known to play an important role in cognitive processes.

Dopamine and noradrenaline are the prominent candidate molecules for mediating the outlined effects of ageing on brain functioning. Both dopaminergic and noradrenergic systems cover widespread connections throughout the brain and are strongly associated with cognitive functioning. Dopamine is mainly synthesised by neurons located in two adjacent midbrain structures: ventral tegmental area (VTA) and substantia nigra pars compacta (SNc; Bissonette and Roesch, 2016), while locus coeruleus (LC), located in the upper dorsolateral part of the brainstem, constitutes the primary source of noradrenaline in the brain (Szabadi, 2013).

Populations of dopaminergic neurons in the midbrain are implicated in reward processing by encoding incentive salience of the stimuli (Berridge and Kringelbach, 2015) and reward prediction error (Schultz, 2016). The research in animal models and humans supports the involvement of the dopaminergic system in cognitive functions such as sustained attention (Bellgrove et al., 2005), working memory (Bäckman and Lyberg, 2013), executive control (Saloner et al., 2020), as well as visuomotor search (Chen et al., 2020). Several resting-state studies have found the functional connectivity of VTA and SNc to be related to aspects of reward processing (Adrián-Ventura et al., 2019; Schmaal et al., 2012), impulsivity (Grimm et al., 2020), interoception (Chong et al., 2017) and gait speed (Karim et al., 2020).

In the course of healthy ageing, the reduction in the number of dopamine receptors and transporters was observed (Karrer et al., 2017) and lifespan alterations in the dopaminergic transmission are quite likely to at least partially mediate the effects of age on different aspects of cognition, proving it to be a solid target for studying the trajectory of human ageing. To our knowledge, only three studies have been published so far regarding age-related changes in the connectivity of main dopaminergic regions. In individuals ranging from 18 to 49 years old, increased age was related to stronger coupling between VTA/SNc and lateral prefrontal cortex, superior temporal and parahippocampal gyri, as well as cerebellum (Manza et al., 2015; Peterson et al., 2017; Zhang et al., 2016). Unlike VTA, SNc was also found to have greater connectivity with angular gyrus (Peterson et al., 2017). The only negative association with age was reported for the coupling between VTA/SNc and somatomotor cortex (Manza et al., 2015; Peterson et al., 2017; Zhang et al., 2016).

The locus coeruleus-noradrenergic system (LC-NE) seems to be especially sensitive to degenerative changes related to ageing, as the LC has been found to be one of the first structures where degenerative changes in Alzheimer’s disease occur (Mravec et al., 2014). Furthermore, the presence of α-synuclein pathology in this structure precedes changes in SNc in Parkinson’s disease (Del Tredici et al., 2002).

Noradrenergic fibres from the LC project to the cerebral cortex, diencephalon, limbic system, cerebellum and through the descending pathway to the spinal cord (Szabadi, 2013). Dense projections from the LC to the fronto-parietal areas and cingulate cortex indicate its role in executive functions, working memory, and attention (Ramos and Arnsten, 2007). Studies with pharmacological modulation of noradrenergic receptors confirmed the role of the NE in the control of working memory in different species, for the α_1_-antagonist phenylephrine, when infused into the dorsolateral prefrontal cortex (DLPFC), was found to impair working memory performance in rats (Arnsten et al., 1999). Meanwhile, administration of α_2_-agonist guanfacine improved performance in the spatial working memory task in rhesus monkeys (Avery et al., 2000) and humans (Jäkälä et al., 1999).

Zhang and colleagues were the first to examine brain functional connectivity of the LC in humans using resting-state fMRI and additionally investigated the effects of ageing on its connectivity patterns (Zhang et al., 2016). They found that age was positively correlated with the strength of connections of the LC to the medial frontal gyrus and cerebellum and negatively correlated with the connectivity to the parahippocampal gyrus and precuneus. Jacobs and others revealed a positive association between the connectivity of the left LC to the left parahippocampal and memory performance among healthy older subjects (Jacobs et al., 2015). Meanwhile, the connectivity between these structures was reduced in patients with mild cognitive impairment (MCI; Jacobs et al., 2015). Two studies revealed nonlinear relationships between the age and LC connectivity. A curvilinear association between the age and functional connectivity of the left LC to the bilateral cortical areas such as temporal pole, posterior middle temporal gyrus angular gyrus, and inferior frontal gyrus, as well as right posterior cingulate cortex was observed by Jacobs and colleagues (Jacobs et al., 2018). Recently, Song, Neal and Lee found a positive quadratic relationship between the age and LC connectivity with sensory regions and a negative quadratic association between the age and LC connectivity with frontal regions (Song et al., 2021).

In this study we focused on dopaminergic and noradrenergic systems considering their important role in cognitive functioning and the evidence of substantial changes in these systems in the course of ageing. With the usage of open-access data, we aimed to investigate whether ageing affects functional connectivity of dopaminergic and noradrenergic systems’ targets over the whole brain. This work extends the previous findings by comparing the resting-state connectivity patterns of VTA, SNc and LC between young (aged 20-35) and older adults (aged 55-80), i.e., beyond the age ranges used in the published papers so far. Secondly, this study aimed to investigate the relationship between behavioural indices of cognitive and motivational processing and the functional connectivity patterns of the primary noradrenergic and dopaminergic nuclei. To the best of our knowledge, no previous works have attempted to find such associations for sustained attention, working memory, visuomotor search and set shifting. Additionally, this is the first study to investigate age-related changes in ReHo within the dopaminergic midbrain. We expected mostly age-related decreases in resting-state functional connectivity of the noradrenergic and dopaminergic seeds observed across multiple cortical and subcortical regions, however, we did not exclude the possibility of increases in functional connectivity of both systems as they have been reported before.

## 2. Materials and methods

### 2.1. Dataset

The anonymised MRI and behavioural data was obtained from the Max Planck Institut Leipzig Mind-Brain-Body Dataset (Babayan et al., 2019). Out of 227 subjects available in the database, only 134 met the initial inclusion criteria, i.e., right-handedness, secondary level education, no psychiatric, neurological or substance abuse history, drug-free. The participants’ age was provided only as a five-year bracket instead of specific values for stronger anonymity protection. Thus, the selected subjects were put into two age groups: 20-35 (N=92) and 55-80 (N=42). All participants from the elderly group matching the inclusion criteria were included in the sample. The size of the younger group was set to match the former, and the subjects were selected randomly. As a result of the fMRI data preprocessing, 5 subjects from the younger group and 15 participants from the elderly group were excluded from the analysis due to: excessive head motion (>fMRI voxel size. i.e., 2.3 mm; 4 younger and 10 elderly), significant brainstem signal loss resulting from susceptibility artefacts (1 younger and 2 elderly), and unsatisfactory brainstem alignment between anatomical and functional data (3 elderly). Thus, the final sample consisted of 37 younger (16 females) and 27 elderly (14 females) subjects. More motion-related dropouts in the older group are typical for fMRI experiments (Madan, 2018).

### 2.2. Behavioural data

The behavioural data chosen for the analyses was associated with several cognitive domains presumably affected by the dopamine and noradrenaline networks activity. Sustained attention was measured as a mean reaction time in the Alertness subtest of the Test of Attentional Performance (TAP) (Zimmermann and Fimm, 2012). Verbal working memory capacity was tested with the Working Memory subtest of TAP (Zimmermann and Fimm, 2012). The mean reaction times of the correct responses and the percentage of correct trials were chosen as variables of interest. The visuomotor capabilities (visual search) and executive functions (set shifting) were investigated using Trail Making Test (TMT) Part A and Part B (Reitan and Wolfson, 2004). The completion time of TMT-A was treated as a measure of visuomotor search capability, while the difference between the completion times of TMT-B and TMT-A was taken into the analyses as the index of set shifting. Lastly, the reward sensitivity was assessed with Reward Responsiveness subscale of the Behavioural Approach System’s (BAS) questionnaire (Carver and White, 1994), with the total score included in the behavioural models. The behavioural paradigms are described in more detail in the Supplementary Data S1.

### 2.3. Behavioural data group-level analysis

The statistical analysis of behavioural data was performed in R (version 4.0.3; R Core Team, 2021). Separate models were created for each measure. They consisted of the behavioural score as the dependent variable, and independent variables age (factor) and sex (covariate). All subjects received the same level of education (i.e.,, secondary), thus it was not included in all the designs throughout the study. The parametric ANCOVA computations were performed with the use of anova function. In the case the data did not adhere to the assumptions of parametric analysis (i.e.,, non-normal distribution of the residuals), a non-parametric rank-based ANCOVA (raov function; Kloke and McKean, 2020) was utilised. The correction for multiple comparisons was achieved by using false discovery rate (FDR). Due to missing behavioural data, one subject (1 elderly female) was excluded from all the TMT-A and TMT-B analyses, and four participants (1 young female, 2 young males, 1 elderly male) were not included in all the BAS Reward Responsiveness models.

### 2.4. MRI data acquisition and preprocessing

The MRI scanning sessions were performed on a 3 Tesla scanner (MAGNETOM Verio, Siemens Healthcare GmbH, Erlangen, Germany) equipped with a 32-channel head coil. High-resolution anatomical images were acquired using T1 MP2RAGE sequence (176 sagittal slices; 1 mm isotropic voxel size; TR = 5000 ms; TE = 2.92 ms; flip angle 1/flip angle 2 = 4°/5°; GRAPPA acceleration factor 3). The T2*-weighted resting-state fMRI data was collected with gradient echo echo planar multiband imaging sequence (64 axial slices acquired in interleaved order; 2.3 mm isotropic voxel size; TR = 1400 ms; TE = 30 ms; flip angle = 69°; multiband acceleration factor = 4). Participants were instructed to stay awake and lie still with their eyes open while looking at a low-contrast fixation cross. The functional data of each subject consisted of 657 volumes, resulting in 15 mins 30 seconds scan duration.

The anatomical data was preprocessed in the Computational Anatomy Toolbox (CAT12; Gaser and Dahnkne, 2016). Each subject’s T1-weighted image was skullstripped and segmented into grey matter (GM), white matter (WM) and cerebrospinal fluid (CSF). The resulting brain image was used for coregistering the functional data, whereas the WM and CSF segments were binarized and resampled to the fMRI resolution for the signal from these tissues to be extracted and used as regressors in the resting-state data denoising.

The functional data preprocessing stream was performed in FSL (fieldmap correction; Jenkinson et al., 2012) and AFNI (all other steps; Cox, 1996). The initial steps included: deleting the first five volumes (to allow for signal equilibration and steady state), despiking, slice-timing, motion (to the median volume) and fieldmap corrections. Resulting images were later coregistered to the skullstripped anatomical data. Subsequently, they were prewhitened, detrended, denoised (14 regressors: 6 original motion parameters; 6 motion parameters derivatives; WM and CSF signal) and band-pass filtered in the 0.009-0.08 Hz range.

### 2.5. Seed regions and whole-brain functional connectivity

The seed regions for resting-state functional connectivity analyses were derived from probabilistic atlases. Due to small volume and close proximity of bilateral VTA, we decided to use a single region of interest (ROI) encompassing the structures on both sides of the brain, based on the work of Murty et al. (2014). In turn, the left and right SNc seeds were taken from the atlas created by Pauli and colleagues (Pauli et al., 2018), which delineated the substantia nigra (SN) subdivisions and thus enabled us to locate the signal of interest more precisely. The left and right LC ROIs were created based on the mask created by Betts and colleagues (Betts et al., 2017).

To reduce the extent of mixing the signal between VTA and bilateral SNc seeds through smoothing, as well as the potential effect of differences in ROI size on observed connectivity patterns, we thresholded the probabilistic ROIs. The arbitrarily-chosen cut-off points were: 0.97 for the VTA seed, and 0.1 for the bilateral SNc seeds. No such thresholding was applied to the bilateral LC seeds. The resulting five ROIs were nonlinearly transformed from MNI space to each subject’s brain image. Figure 1 shows the location of the seeds in an example participant of our study. Across the cohort, the average ROI volume was: 349.61 (± 53.94) mm^3^ for VTA, 278.89 (± 45.28) mm^3^ for the left SNc, 288.40 (± 40.48) mm^3^ for the right SNc, 89.95 (± 28.21) mm^3^ for the left LC, and 90.24 (± 29.32) mm^3^ for the right LC.

**Figure 1.**
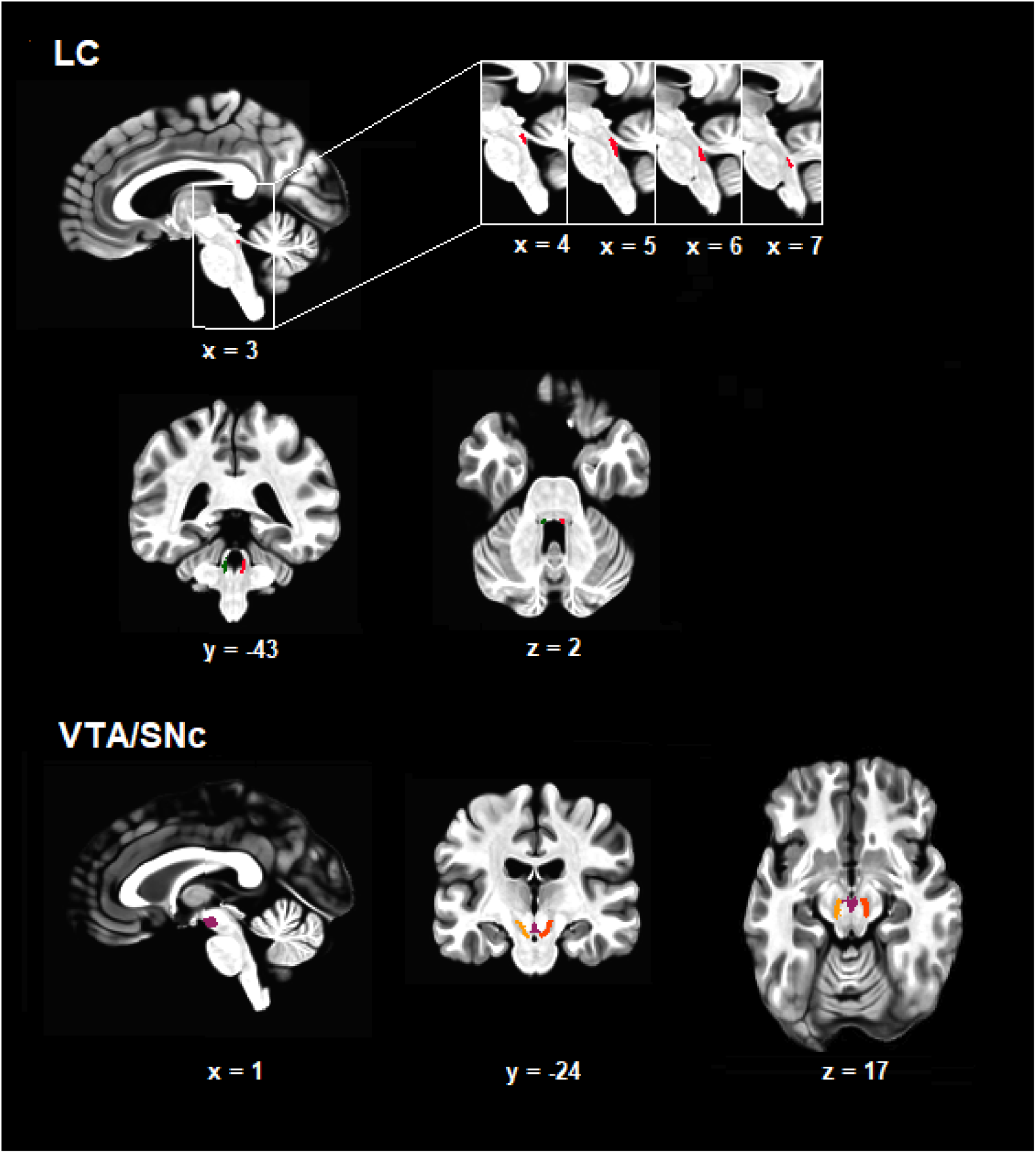
Location of regions of interest (ROIs) in an example subject of the current study. The panel above shows the location of the left (dark green) and right (red) locus coeruleus (LC). The panel below shows the location of bilateral ventral tegmental area (VTA; purple), as well as the left (orange) and right (red) substantia nigra pars compacta (SNc).

**Figure 2.**
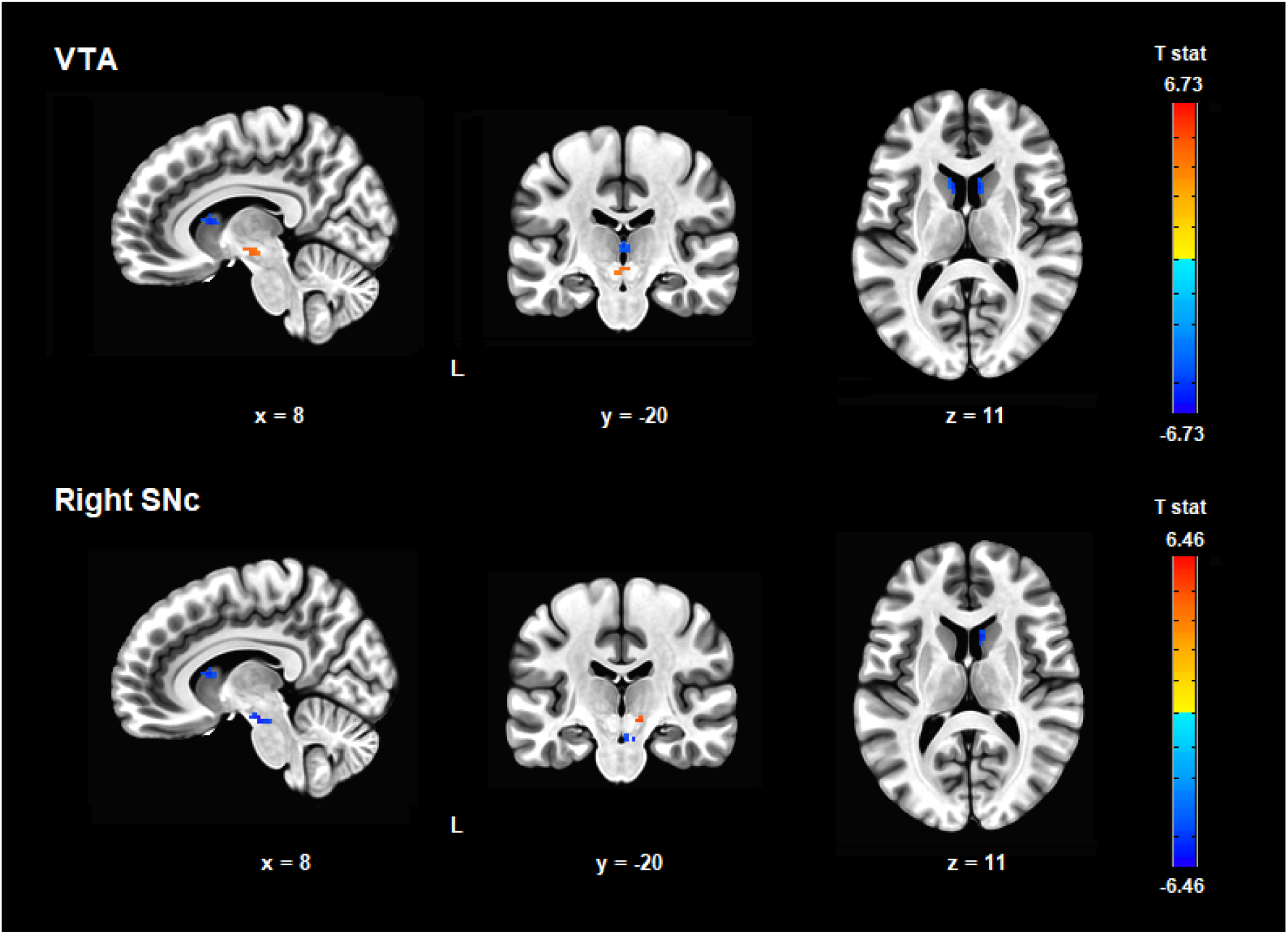
Areas of increased (warm colours) and decreased (cold colours) resting-state functional connectivity with VTA and the right SNc in the elderly group.

**Figure 3.**
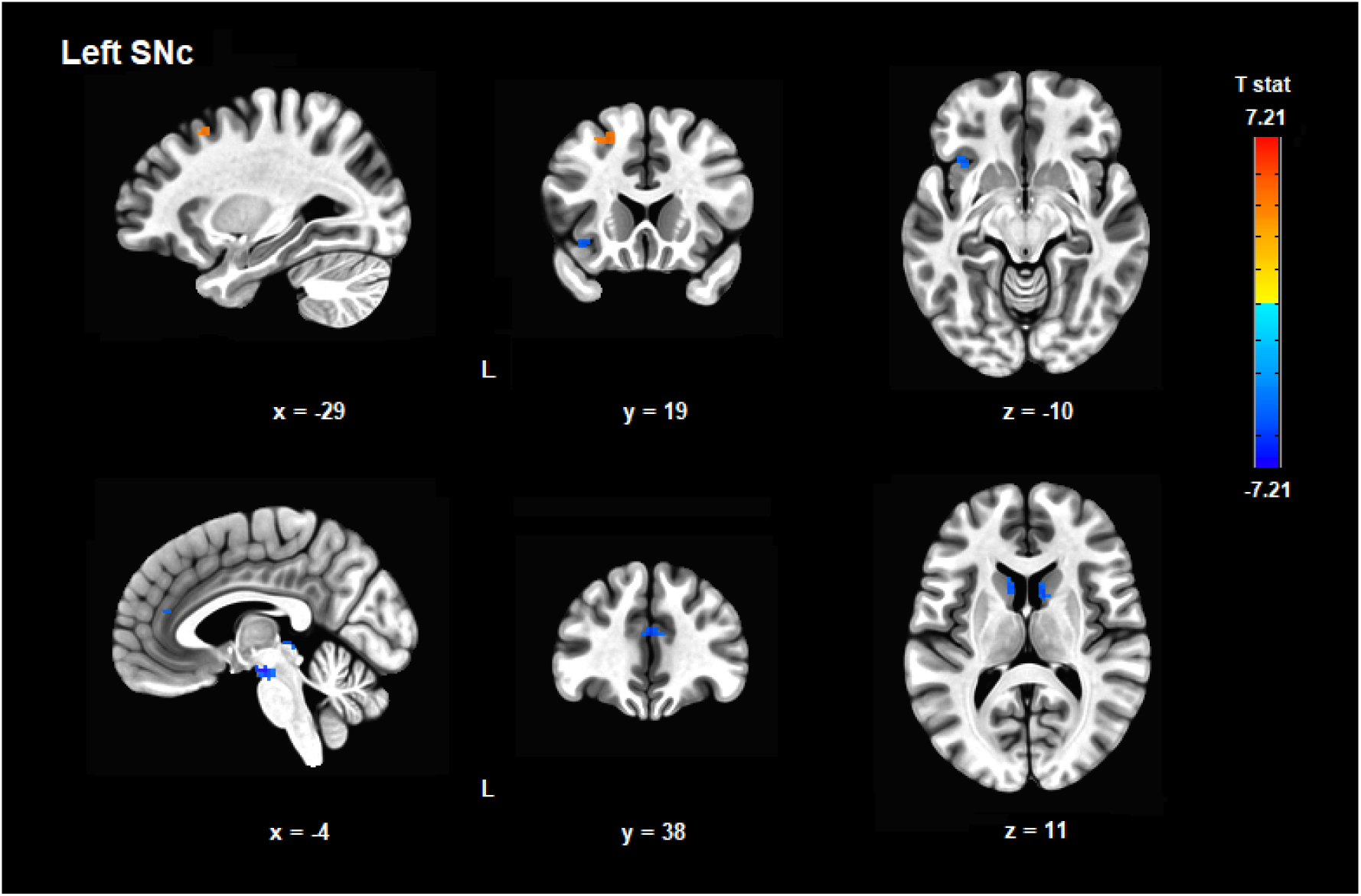
Areas of increased (warm colours) and decreased (cold colours) resting-state functional connectivity with the left SNc in the elderly group.

**Figure 4.**
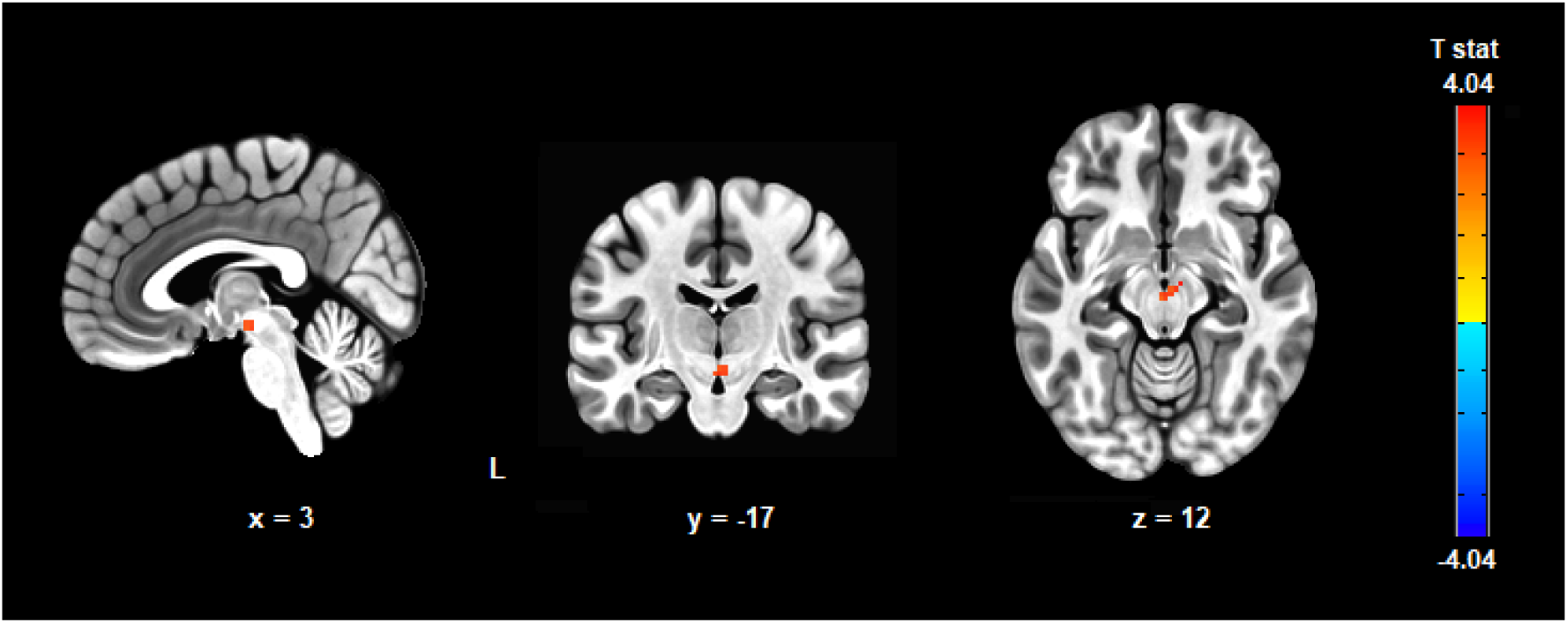
The elderly group was found to have increased regional homogeneity (ReHo) within VTA.

Prior to performing whole-brain seed-based functional connectivity calculations, the preprocessed resting-state volumes were spatially smoothed with a Gaussian kernel of 4 mm FWHM for VTA and bilateral SNc, and with a Gaussian filter of 3 mm FWHM for bilateral LC. Subsequently, the seed time courses of the BOLD signal were produced by averaging BOLD time-series from all voxels in each seed. The correlations were computed between these mean time courses of every ROI and all other voxels in the brain. To normalise the data distribution, the Pearson correlation coefficients r were converted to z-scores using Fisher’s z transform. The resulting maps were normalised to the MNI space to allow group-level analysis.

### 2.6. Seed-based functional connectivity group-level analysis

All statistical analyses of the seed-based whole-brain functional connectivity data were performed using AFNI’s 3dMVM program (Chen et al., 2014). The primary model consisted of age group as the factor, with sex as a covariate. The following comparisons were made: (i) one sample t-tests of VTA, the left and right SNc, and the left and right LC functional connectivity; (ii) paired t-tests comparing the connectivity patterns of the bilateral SNc and LC in the whole sample; (iii) independent samples (old versus young) t-tests of each seed’s whole-brain functional connectivity.

To assess the resting-state functional connectivity patterns associated with behavioural results, we additionally performed the multiple regression computations with resting-state functional connectivity maps as dependent variables, behavioural measures as the predictors, and sex as a confound, separately for each of the five behavioural domains (i.e., sustained attention, visuomotor search, set shifting, working memory and reward responsiveness) and each of the five seeds. As the results were available in the form of F-statistics, which precludes assessing the direction of effects, mean Z values from the suprathreshold clusters were extracted and subsequently correlated with behavioural scores using R (version 4.0.3).

For the one sample t-tests of seed-based functional connectivity, the correction for multiple comparisons was achieved through cluster-level family-wise error rate (FWE). In the case of all other analyses, voxel-level FDR was applied. The respective FDR cluster-forming thresholds for each analysis are provided in the Results section. The decision to use FWE for one sample t-tests of seed-based functional connectivity was guided by the fact that resting-state coupling patterns of VTA, SNc and LC have already been established in the literature, and this analysis served as a means of quality control for seed location. On the other hand, using FDR correction in the remaining analyses was meant to increase their statistical power.

### 2.7. Regional homogeneity calculation and group-level analysis

To assess whether ageing was also associated with changes in the local resting-state functional connectivity within the dopaminergic regions, ReHo analysis was performed (Zang et al., 2004). The ReHo measures the similarity of time-courses of the neighbouring voxels (7 in the case of our study) in a voxel-wise manner using Kendall’s coefficient of concordance (KCC). Due to the small volume of LC ROIs in our study, their oblong shape and relatively large distance between the left and right ROI, we decided to leave out the ReHo calculation for the noradrenergic seeds.

ReHo calculations were performed for the preprocessed resting-state data with the usage of AFNI’s 3dReHo program. To standardise the scores of each participant, the ReHo value of each voxel was divided by the respective brain-wise mean ReHo. No spatial smoothing was applied. The resulting images were transformed into the MNI space to allow a group-level comparison. The statistical analysis was done with the small volume correction (SVC; i.e., within a mask consisting solely of the dopaminergic seeds) using AFNI’s 3dMVM program (Chen et al., 2014). The model consisted of age (factor) and sex (covariate). The correction for multiple comparisons was performed with voxel-level FDR.

## 3. Results

### 3.1. Whole-brain functional connectivity of VTA, SNc and LC

Whole-brain functional connectivity of examined seeds was assessed with cluster-level FWE (p < 0.05) following cluster-forming threshold at uncorrected p < 0.001. The connectivity patterns are depicted in Supplementary Figures 1-5. All dopaminergic ROIs had positive connectivity to bilateral cingulate, dorsomedial prefrontal and lateral orbitofrontal cortex, inferior frontal gyrus pars opercularis, parieto-occipital sulcus, the medial temporal cortices, insula, caudate nucleus, putamen, nucleus accumbens and globus pallidus, thalamus, pons, midbrain and cerebellum. Similarly, they shared negative connectivity to bilateral motor and somatosensory cortices, as well as parts of the superior and inferior parietal lobes. All seeds were furthermore negatively connected to the visual areas, with the positive coupling between the right SNc and bilateral primary visual cortex being the only exception.

Both left and right LC showed positive connectivity to bilateral anterior cingulate, thalamus, hippocampus, cerebellum and brainstem and negative connectivity to bilateral primary motor and somatosensory cortices, precuneus, superior parietal lobules, as well as the right inferior parietal lobule and the right superior frontal gyrus. The right, but not left, LC was positively coupled with the bilateral hippocampus and negatively coupled with the left superior frontal gyrus. The left LC was negatively connected to bilateral visual cortices, whereas in the right LC the relationship was present only for the ipsilateral regions.

Despite the observed differences in one sample t-tests results, a direct comparison of the connectivity strength between the left and right LC yielded no significant results (voxel-level FDR < 0.05; minimal cluster size k = 10 voxels). In turn, contrasting the coupling patterns of the left and right SNc revealed that each seed showed greater coupling with cerebral cortex and midbrain regions located ipsilaterally to the given ROI (voxel-level FDR < 0.05; k = 40). The opposite connectivity pattern was observed for the cerebellum as the left SNc was more strongly connected to the right cerebellar hemisphere. The results are presented in more detail in Supplementary Figure 6 and Supplementary Table 1.

### 3.2. The impact of age on connectivity patterns

No effect of age on the connectivity of the left or right LC was found (voxel-level FDR < 0.05; k = 10). In turn, a number of differences was observed for the dopaminergic seeds (voxel-level FDR < 0.05; k = 10). The older group had decreased functional connectivity between VTA and bilateral caudate nuclei and mediodorsal thalamus, between the left SNc and bilateral caudate nuclei and anterior cingulate cortex (ACC), as well as the left superior colliculus and insula, and between the right SNc and right caudate nucleus. Increased age was also associated with greater signal correlation between the left SNc and the ipsilateral DLPFC. The difference in age was further linked to different patterns of connectivity to the midbrain structures. In older adults, VTA had higher resting-state connectivity to bilateral red nucleus, while the right SNc was more strongly connected to the midbrain area adjacent to the unilateral red nucleus. At the same time, bilateral SNc seeds showed a decreased extent of connectivity to VTA and neighbouring voxels.

### 3.3. Age-related differences in regional homogeneity of the dopaminergic seeds

The analysis revealed that the older group had increased ReHo within VTA (voxel-level FDR < 0.05; k = 7). Suprathreshold statistics were found for a cluster consisting of 12 out of 105 voxels from the SVC mask (MNI: x = 1, y = -17, z = -12; peak voxel T-stat = 3.67).

### 3.4. Age-related differences in behavioural measures

The older group was characterised by decreased sustained attention (longer reaction times in TAP Alertness; FDR = 0.002), visuomotor search capability (longer completion time of TMT-A; FDR < 0.001) and set shifting (greater difference between TMT-B and TMT-A completion times; FDR < 0.001). Interestingly, while the older participants performed worse in the TAP Working Memory task (i.e., fewer correct responses; FDR = 0.020), the reaction times of the successful trials did not differ significantly between the groups (FDR = 0.081). Additionally, the older and younger subjects scored alike in BAS Reward Responsiveness (FDR = 0.306). The results are described in more detail in the Supplementary Table 2.

### 3.5. Functional connectivity associated with behavioural results

The only significant associations between resting-state functional connectivity and behavioural measures were found for the right SNc and the TMT-A visuomotor task (voxel-level FDR < 0.05; k = 10). The connectivity strength of the right SNc with both the left angular gyrus (R = 0.67) and the right lateral orbitofrontal cortex (R = 0.64) was positively correlated with the completion time of the task, i.e., the greater the coupling, the worse the performance. The results are shown in more detail in Table 2 and Figure 5. The correlation plots of the TMT-A completion time and connectivity strength between the brain areas are shown in Supplementary Figures 7 and 8.

**Table 1.** Age-related differences in resting-state functional connectivity (voxel-level FDR < 0.05 and cluster extent of minimum 10 voxels).

| **MNI** | **Location** | **Voxels** | **Directionality** | **T-stat** |
| --- | --- | --- | --- | --- |
| *VTA* |  |  |  |  |
| 5, -15, 10 | B red nucleus | 31 | Old > Young | 6.72 |
| 8, 12, 8 | R caudate nucleus | 16 | Young > Old | 5.82 |
| -9, 12, 11 | L caudate nucleus | 15 | Young > Old | 5.58 |
| 1, -20, 6 | B mediodorsal thalamus | 10 | Young > Old | 5.32 |
| *L SNc* |  |  |  |  |
| -4, -15, -17 | B ventral tegmental area | 50 | Young > Old | 7.20 |
| 10, 10, 11 | R caudate nucleus | 23 | Young > Old | 5.52 |
| -9, 12, 11 | L caudate nucleus | 20 | Young > Old | 5.33 |
| 1, 38, 18 | B anterior cingulate cortex | 20 | Young > Old | 5.76 |
| -29, 17, 50 | L dorsolateral prefrontal cortex | 20 | Old > Young | 4.88 |
| -2, -31, -1 | L superior colliculus | 18 | Young > Old | 5.69 |
| -36, 19, -10 | L insula | 12 | Young > Old | 4.98 |
| *R SNc* |  |  |  |  |
| 5, -15, -17 | B ventral tegmental area | 24 | Young > Old | 6.46 |
| 8, 15, 11 | R caudate nucleus | 12 | Young > Old | 5.57 |
| 12, -20, -8 | R midbrain | 12 | Old > Young | 5.50 |

**Table 2.** Resting-state functional connectivity patterns of dopaminergic seeds correlated with behavioural performance (voxel-level FDR < 0.05 and cluster extent of minimum 10 voxels).

| Seed | MNI | Location | Voxels | Pearson's correlation [r] | F-stat |
| --- | --- | --- | --- | --- | --- |
| *TMT-A* |  |  |  |  |  |
| R SNc | -36, -61, 36 | L angular gyrus | 27 | 0.67 | 28.28 |
| R SNc | 37, 42, -15 | R orbitofrontal cortex | 14 | 0.64 | 37.69 |

**Figure 5.**
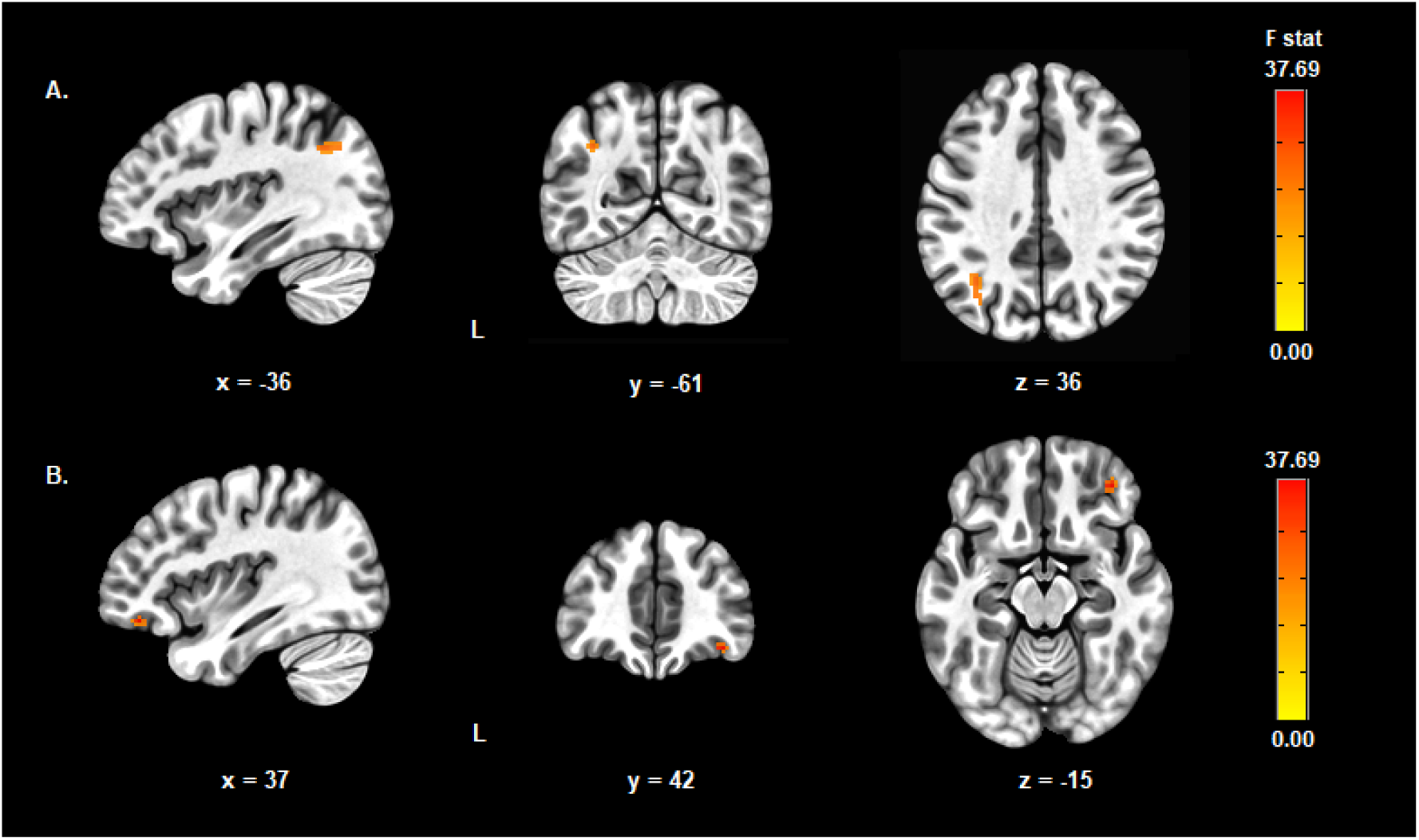
The functional connectivity of the right SNc with (A) the left angular gyrus and (B) the right lateral orbitofrontal cortex was found to be positively correlated with the completion time of the TMT-A task probing visuomotor search capabilities.

## 4. Discussion

### 4.1. Functional connectivity patterns of VTA, SNc and LC

The dopaminergic connectivity patterns in our study considerably overlap with those reported in earlier papers (Manza et al., 2015; Peterson et al., 2017; Zhang et al., 2016). The obtained pattern of resting state functional connectivity of the LC, comprising the widespread connections to the cortical and cerebellar areas, is also consistent with previous studies conducted both on animal and human models (Zhang et al., 2016; Liebe et al., 2020). To the best of our knowledge, this is the first study to show the differences in the functional connectivity between the left and right SNc. The observed coupling pattern, i.e., greater connectivity with ipsilateral subcortical and cortical regions along with stronger correlation with the contralateral cerebellum, is in line with the results of anatomical studies on rodents, primates and humans (Molochnikov and Cohen, 2014; Meola et al., 2016). Considering the strong laterality of symptoms at the onset of Parkinson’s disease, treating right and left SNc as separate seed regions could hold some potential in clinical MRI applications (Heinrichs-Graham et al., 2017).

### 4.2. Age-related differences in dopaminergic long-range connectivity

Our study is the first one to show that healthy ageing is associated with decreases in functional connectivity between VTA/SNc and several key telencephalic recipients of dopaminergic innervation, namely ACC, caudate nucleus, and insula. Ageing has been associated with the loss of dopamine transporters in the caudate nucleus and dopaminergic receptors in all three regions (Karrer et al., 2017). Furthermore, administration of L-DOPA, a precursor of dopamine synthesis, was found to increase the VTA/SNc functional connectivity with insula (Grimm et al., 2020). Additionally, the finding of increased coupling of the left SNc with the left DLPFC in older adults corresponds well with an earlier report showing the connectivity between VTA/SNc and lateral prefrontal cortex increased with age (18-49 years old; Manza et al., 2015).

ACC is involved in the action-outcome learning, and one of the pathways through which it guides the selection of future behaviour leads to the DLPFC (Der-Avakian and Markou, 2012). Older adults perform worse in tasks requiring intentional motivated learning with delayed feedback and probabilistic reward learning, showing manifest changes in the associations between taken actions and resulting outcomes, unless these two are paired in a constant fashion and a short time interval (Geddes et al., 2018; Samanez-Larkin et al., 2014). One of these studies found older subjects to have decreased correlation between ACC activity and prediction errors, which may serve as a mechanism responsible for deteriorated action-outcome learning and the reduced functional connectivity observed in our study (Samanez-Larkin et al., 2014).

Caudate, insula and mediodorsal thalamus, where we also found age-related functional connectivity decrease, are all important hubs for integrating/relaying information from multiple neural systems, including emotional, cognitive, sensory and motor circuits (Haber, 2014; Uddin et al., 2017; Mitchell and Chakraborty, 2013). Due to these neural connections, the alterations in functional connectivity with these structures may constitute a correlate of behavioural deterioration in more than one specific domain. Together with the decreased coupling between the left SNc and ipsilateral superior colliculus, which plays a major role in fast processing of visual stimuli and is essential for assigning attentional salience (Comoli et al., 2003), they could contribute to the attentional deficits of the elderly such as suppressed ability to ignore task-irrelevant stimuli (as reviewed in Swirsky and Spaniol, 2019). On the other hand, the decreased connectivity of the dopaminergic seeds to the caudate nucleus could also result in age-related psychomotor slowing. Indeed, decreased coupling within the basal ganglia system was associated with slower gait (Karim et al., 2020). This mechanism could act in convergence with the observed in our study increased coupling of VTA/SNc to the red nucleus, which is known to exert a modulatory effect on cortico-striatal networks and has been implicated in motor functions (Bostan and Strick, 2018; Meola et al., 2016). All in all, these two findings of ours complement previous ageing studies which reported differences in VTA/SNc connectivity with somatomotor cortex and cerebellum, further supporting the role of dopaminergic dysfunction in motor deterioration in older adults (Manza et al., 2015; Peterson et al., 2017).

### 4.3. Age-associated changes in local dopaminergic connectivity

Our analyses revealed age-related changes in the functional organisation within the dopaminergic midbrain. In older adults bilateral SNc seeds were less strongly coupled with VTA. Additionally, the same group was characterised by greater ReHo in VTA. Overall, our study is the first one to indicate that ageing is associated with differences in connectivity between dopaminergic areas, as well as local synchronisation within them. Both dopaminergic structures are a part of the striato-midbrain system, which acts as an interface for relaying information from limbic to cognitive to motor circuits (Haber, 2014). The decrease of functional connectivity between SNc and VTA is in line with a vast body of literature showing that ageing is associated with a decreased strength of coupling within resting-state functional systems (Jockwitz and Caspers, 2021). The local synchronisation, measured with ReHo, has been shown to vary across the brain, with lower values being characteristic for areas with complex functions, presumably stemming from more sophisticated underlying activity (Song et al., 2014). As VTA is known to be less vulnerable for age-related pathological degeneration than SNc, our results might be a marker of the differential ageing trajectories and diminished communication between these two dopaminergic regions (Fu et al., 2016).

### 4.4. Ageing affects several cognitive domains but not reward processing

In line with the previously published research, we found age-related decreases in the following domains: sustained attention (Vallesi et al., 2021), visuomotor search and set shifting (St-Hilaire et al., 2018), as well as working memory accuracy (Bopp and Verhaeghen, 2020). Surprisingly, the group difference in the mean correct response time in the working memory task only approached the significance level (FDR = 0.081). A recently published meta-analysis investigating the effects of age on the performance in the 2-back task, analogous to the TAP Working Memory, showed significant negative effect of age on both accuracy and overall mean response times (Bopp and Verhaeghen, 2020), with the effect size larger for the latter measure. While one key difference between our study and the meta-analysis lies in the use of mean response times for correct versus all responses, we believe that the lack of significant difference in our case stems from inclusion of insufficient number of subjects as meta-analyses are known to have increased statistical power compared to individual investigations. Similar levels of reward responsiveness in the two groups are also in line with a vast body of research showing that value encoding of rewarding outcomes remains intact in the elderly individuals (Samanez-Larkin and Knutson, 2015).

### 4.5. Functional connectivity of the right SNc predicts visuomotor functions

Our analyses revealed that the connectivity strength of the right SNc with both the left angular gyrus and the right lateral orbitofrontal cortex was positively correlated with the completion time of the TMT-A task probing visuomotor capabilities, i.e., the greater the coupling, the worse the performance. The effect sizes of these associations (R = 0.64-0.67) should be, however, treated with caution as circular analyses are known to greatly overestimate the true effect strengths (Kriegeskorte et al., 2010). According to the Neurosynth database, the location of the angular gyrus cluster maps onto the edge of the medial frontoparietal (default mode) network, which is expected not to have positive correlation with areas involved in visuomotor search (Yarkoni et al., 2011). Lateral orbitofrontal cortex, where the other cluster was found, and adjacent parts of the inferior frontal gyrus are known to be involved in the stop-related processing (Deng et al., 2017). They inhibit movement through their influence on the indirect or hyperdirect basal ganglia pathways (Aron and Poldrack, 2006). As elevated dopaminergic transmission is known to promote movement (Minassian et al., 2016), the negative correlation between the right SNc - the right lateral orbitofrontal cortex connectivity and the task performance indicates that anti-correlation between these two structures could facilitate the selection of appropriate motor behaviour, enhancing visuomotor search abilities. Interestingly, the part of the lateral orbitofrontal cortex associated with the performance in the task was different from the one identified to have positive coupling with the right SNc in one sample t-test. This could indicate the existence of subdivisions within the lateral orbitofrontal cortex differentially associated with the dopaminergic transmission.

### 4.6. Age-related connectivity patterns of VTA/SNc are not reflected in changes in cognitive functions

As the age and behavioural measures were not independent, the analyses investigating resting-state substrates associated with behavioural outcome were conducted without accounting for the age of participants to compare the connectivity patterns associated with task performance and ageing. Despite finding age-related VTA/SNc connectivity differences in a number of structures implicated in the studied behavioural paradigms, none of these were replicated in association with the task performance. To the best of our knowledge, only one study reported correlations between cognitive functions tested here (reward sensitivity) and resting-state connectivity of the dopaminergic seeds (Adrián-Ventura et al., 2019). Compared to our report, this work deployed a larger cohort (89 individuals) and analysed the data in the ROI-to-ROI fashion, greatly increasing its power.

### 4.7 No evidence for LC connectivity changes related to ageing and behaviour

This study did not show any significant age- and behaviour-related findings for the LC connectivity, thus failing to reproduce earlier reports, most probably due to the insufficient number of subjects per group. It was shown that the relationship between age and LC connectivity may be more complex than just linear decrease (Jacobs et al., 2018; I. Song et al., 2021). Different results might have been obtained if the older group was divided into middle-aged adults (55-65) and older (65-80) adults. However, the small sample size of the elderly group and the inequality in the number of subjects per age range hindered more specific group assignments. The lack of significant age differences in LC connectivity precluded the comparison of regions affected by ageing between the dopaminergic and noradrenergic system. We also did not find any correlation between the LC resting-state functional connectivity and outcomes of behavioural tests.

### 4.8. Limitations of the study

There are several limitations to this study. As outlined above, larger sample size would have increased the statistical power of the analyses, making it better suited to detect more subtle differences in connectivity patterns. Furthermore, knowing the precise age of participants would have enabled investigation of the linear relationship between the age and resting-state connectivity. Future longitudinal studies are warranted to confirm the cross-sectional age-related differences in the connectivity of VTA and SNc reported in this study.

Taking into account the distinct structural connectivity patterns of SNc subdivisions (Zhang et al., 2017), it would be beneficial to study whether their tractography differences are indeed reflected in resting-state connectivity and behavioural measures these coupling patterns are correlated with. Preferably, the location and delineation between VTA and SNc subdivisions should be achieved in a data-driven manner rather than through the use of atlases like those deployed in this study. Single-subject level localisation with the usage of neuromelanin-sensitive imaging should be used in the future studies to account for interindividual variability in localisation of the SNc and LC. Additionally, this method would enable comparison of the intensity of neuromelanin signal with cognitive functions measured by behavioural tests.

Last but not least, since the LC is known to be involved in the control of the cerebral blood flow and neurovascular coupling, it could be also useful to include in the analyses the measures of the blood flow and pulse.

## 5. Conclusions

Our findings show age-related decrease in VTA/SNc long range connectivity with caudate, mediodorsal thalamus, ACC and insula. Additionally, ageing impacts local connectivity within dopaminergic regions observed as increased VTA ReHo and decreased VTA-SNc coupling. Furthermore, this study shows link between the functional connectivity of SNc and visuomotor performance, as well as underlines the functional differences between bilateral SNc. In conclusion, our study shows that ageing is associated with changes in local connectivity within dopaminergic centres, as well as their coupling with both cortical and subcortical brain regions.

## Data Availability Statement

This work was performed using a widely available open dataset (Babayan et al., 2019).

## Ethics Statement

The following study was not subjected to an ethical committee approval due to being performed on a widely available and fully anonymised dataset.

## Author Contributions

**M.R.Z**.: Conceptualisation, Methodology, Validation, Formal Analysis, Data Curation, Writing - Original Draft, Project administration. **W.F**.: Conceptualisation, Methodology, Validation, Formal Analysis, Data Curation, Writing - Original Draft, Visualisation. **M.B**.: Conceptualisation, Methodology, Writing - Review & Editing, Supervision.

## Supplementary Material

The following Supplementary Material is available alongside the manuscript: Supplementary Data S1: Behavioural tasks description; Supplementary Figure S1: The functional connectivity patterns of bilateral VTA; Supplementary Figure S2: The functional connectivity patterns of the left SNc; Supplementary Figure S3: The functional connectivity patterns of the right SNc; Supplementary Figure S4: The functional connectivity patterns of the left LC; Supplementary Figure S5: The functional connectivity patterns of the right LC; Supplementary Figure S6: Brain areas with differential functional connectivity with the left and right SNc; Supplementary Figure S7: Correlation plot of the TMT-A task completion time and resting-state functional connectivity between the right SNc and the left angular gyrus; Supplementary Figure S8: Correlation plot of the TMT-A task completion time and resting-state functional connectivity between the right SNc and the right lateral orbitofrontal cortex; Supplementary Table S1: The differences in whole-brain resting-state functional connectivity of the left and the right SNc; Supplementary Table S2: The comparison of the behavioural performance between the groups.

