## Supplementary Data S1 for "Influence of age and cognitive performance on resting-state functional connectivity of dopaminergic and noradrenergic centres"

**Supplementary Data S1: Behavioural tasks description**

In the Alertness subtest of Attentional Performance (TAP) (Zimmermann and Fimm, 2012), participants had to press a button when a visual stimulus (cross) appeared on screen. The paradigm was administered with two conditions (with or without audio signal before stimuli). Two repetitions of each condition (1. and 4. without audio signal; 2. and 3. with audio signal), each containing 20 stimuli, were performed. The mean reaction times in both conditions were calculated separately, and their average was the basis of the behavioural analyses. In the Working Memory subtest of TAP (Zimmermann and Fimm, 2012), numbers from 1 to 9 appeared serially on the screen, and the task was to press a button when the number on the screen was equal to the second last number. In the Trail Making Test (TMT) Part A, numbers from 1 to 25 were presented and participants had to connect them in the ascending order (Reitan and Wolfson, 2004). In the part B, numbers from 1 to 13 and letters from A to L were shown, and subjects were tasked with connecting numbers and letters in an alternating and ascending order (i.e., 1-A-2-B-3-C, etc.). Last but not least, the Reward Responsiveness subscale of the Behavioural Approach System's (BAS) questionnaire consisted of 5 items about anticipation or occurrence of reward, each rated from 1 (strongly disagree) to 4 (strongly agree) (Carver and White, 1994).

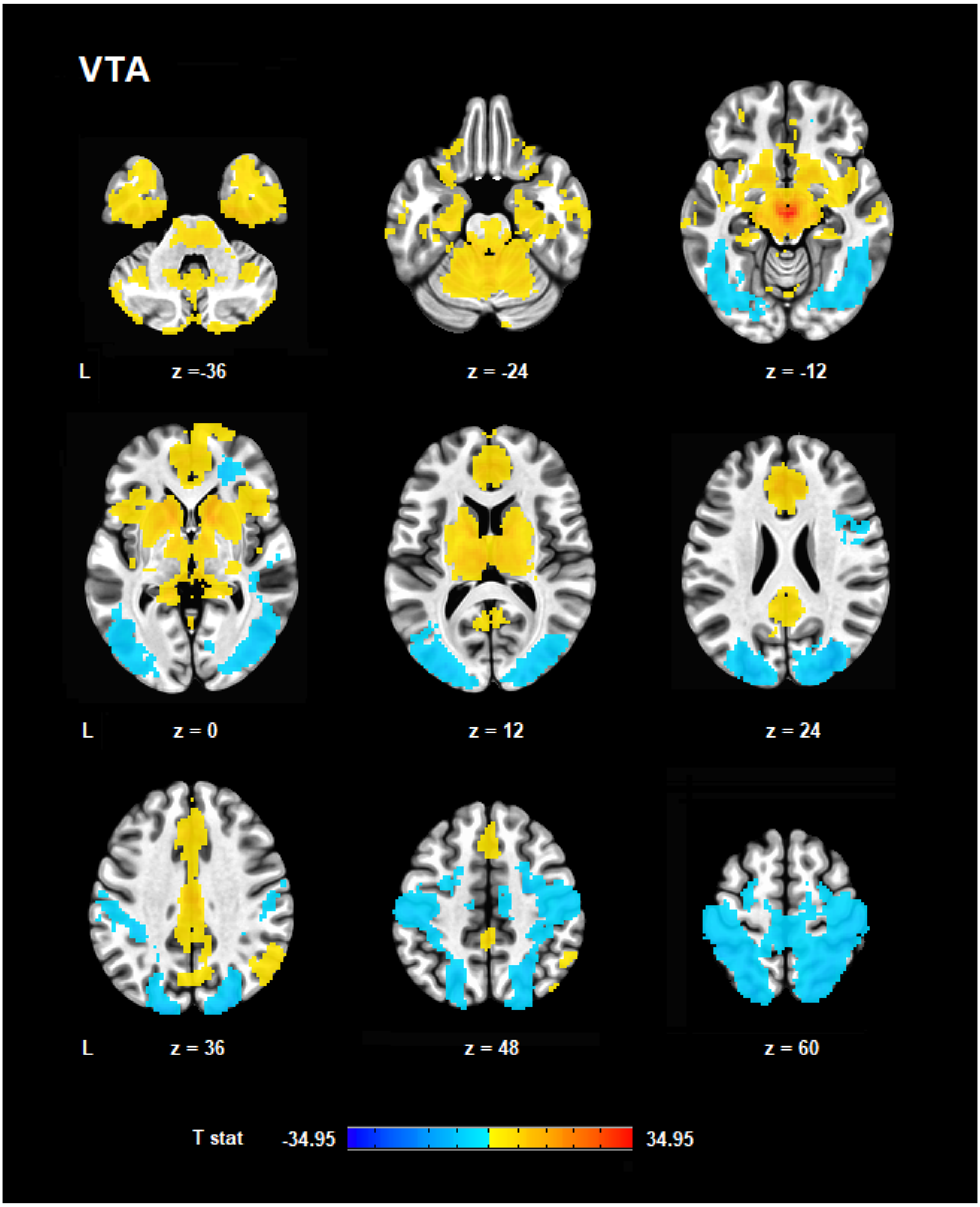


**Supplementary Figure S1.** The functional connectivity patterns of bilateral ventral tegmental area (VTA) assessed with cluster-level FWE ($p < 0.05$) following cluster-forming threshold at uncorrected $p < 0.001$. Warm and cold colours indicate, respectively, positive and negative correlations.

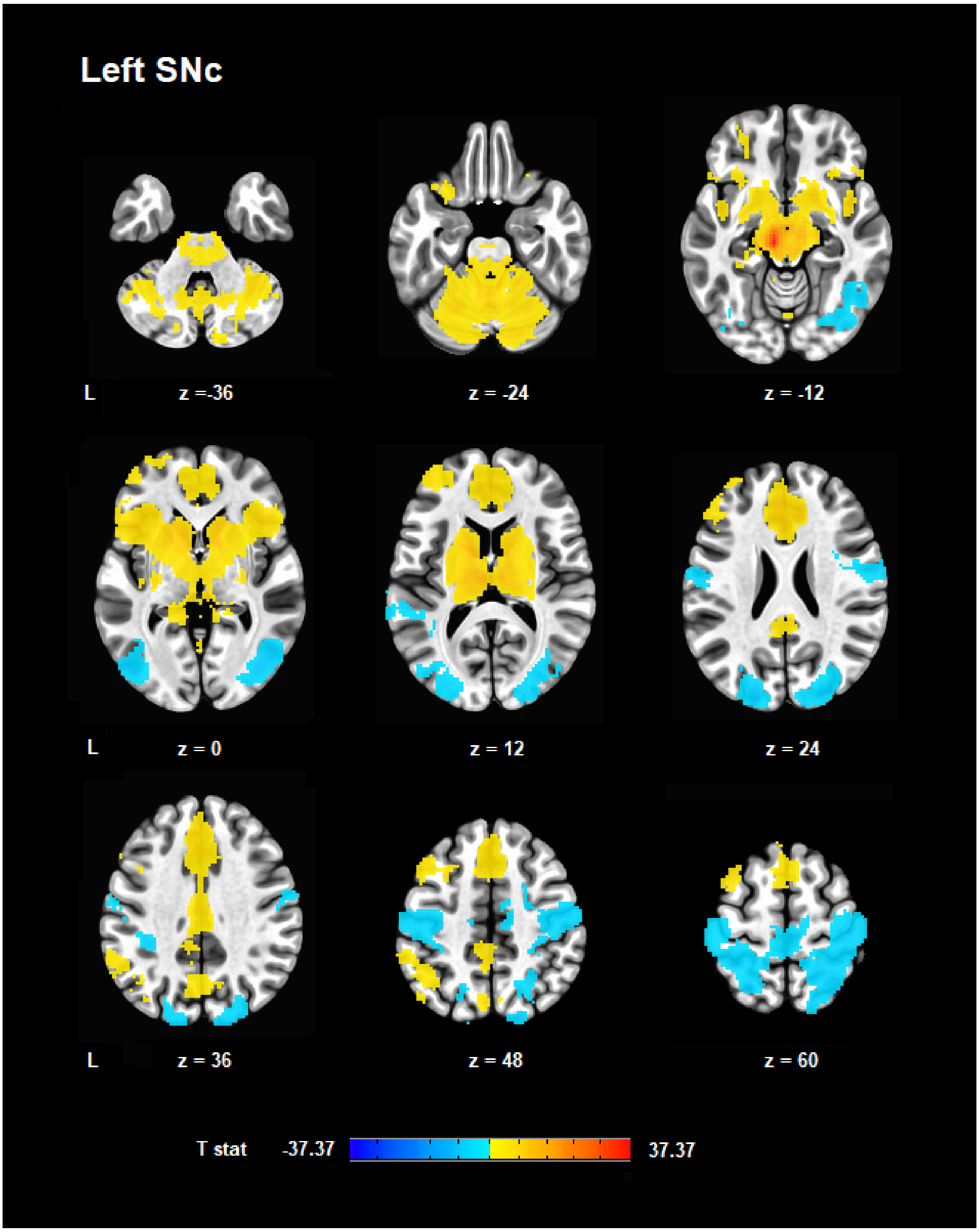


**Supplementary Figure S2.** The functional connectivity patterns of the left substantia nigra pars compacta (SNc) assessed with cluster-level FWE ($p < 0.05$) following cluster-forming threshold at uncorrected $p < 0.001$. Warm and cold colours indicate, respectively, positive and negative correlations.

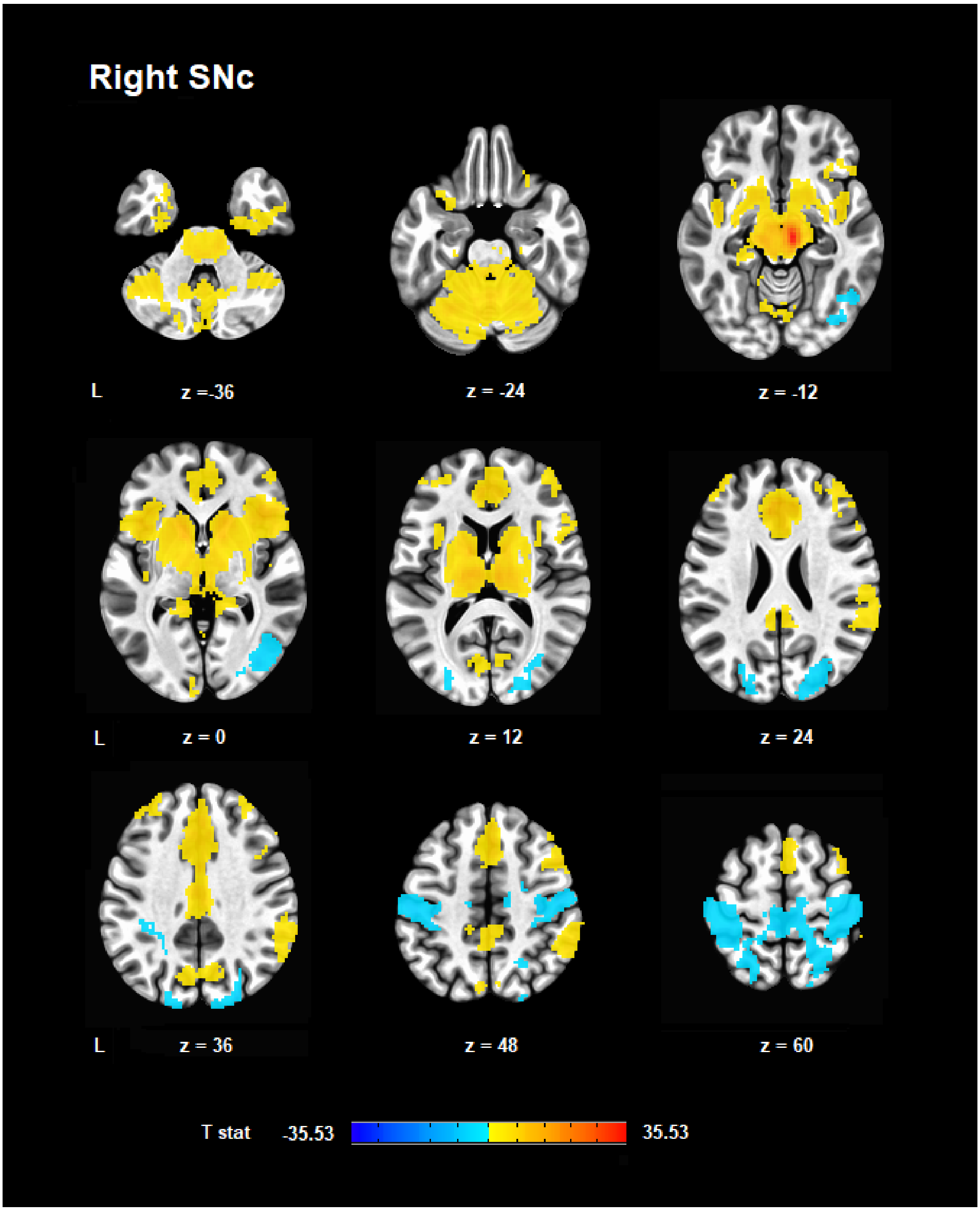


**Supplementary Figure S3.** The functional connectivity patterns of the right SNc assessed with cluster-level FWE ($p < 0.05$) following cluster-forming threshold at uncorrected $p < 0.001$. Warm and cold colours indicate, respectively, positive and negative correlations.

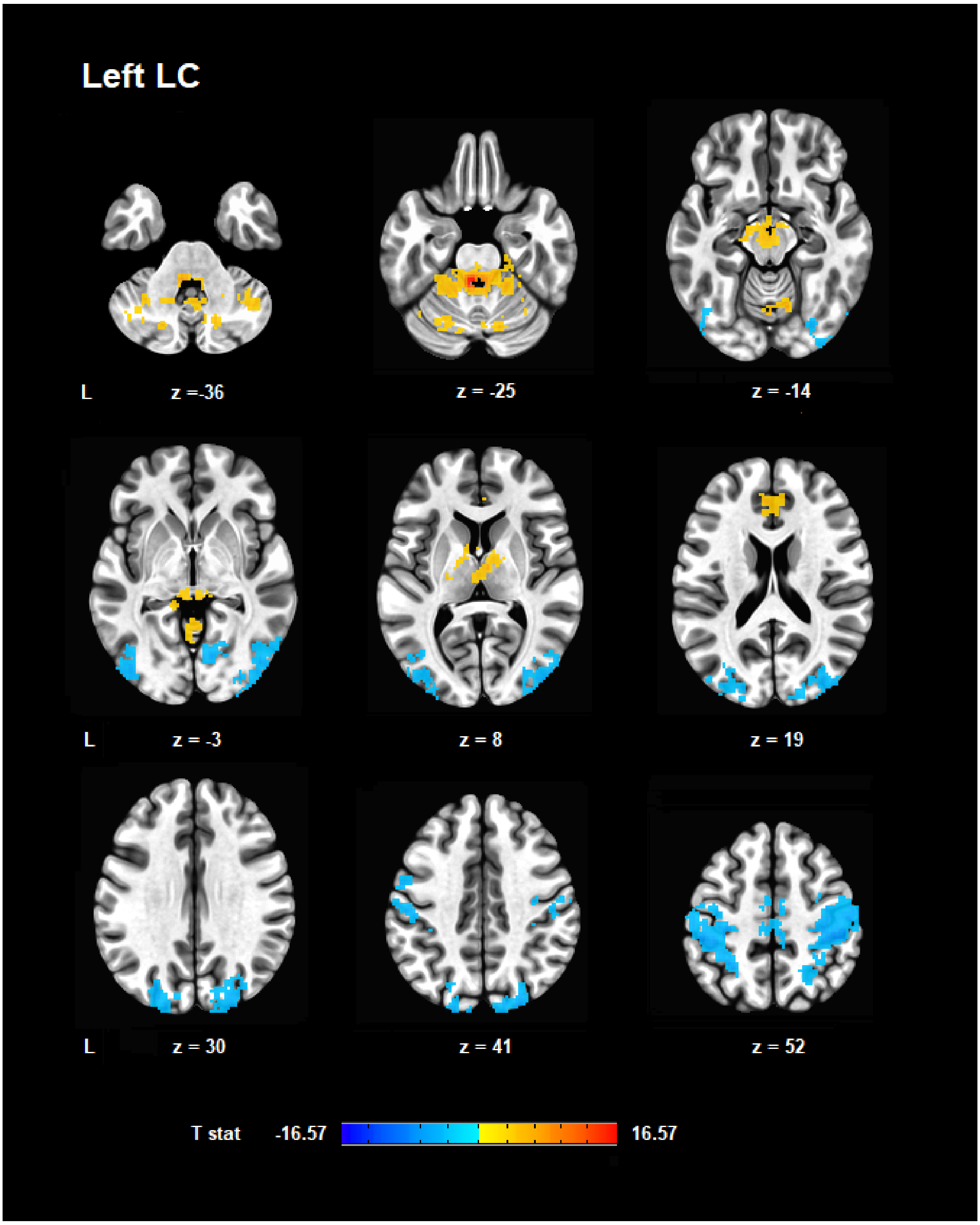


**Supplementary Figure S4.** The functional connectivity patterns of the left locus coeruleus (LC) assessed with cluster-level FWE ($p < 0.05$) following cluster-forming threshold at uncorrected $p < 0.001$. Warm and cold colours indicate, respectively, positive and negative correlations.

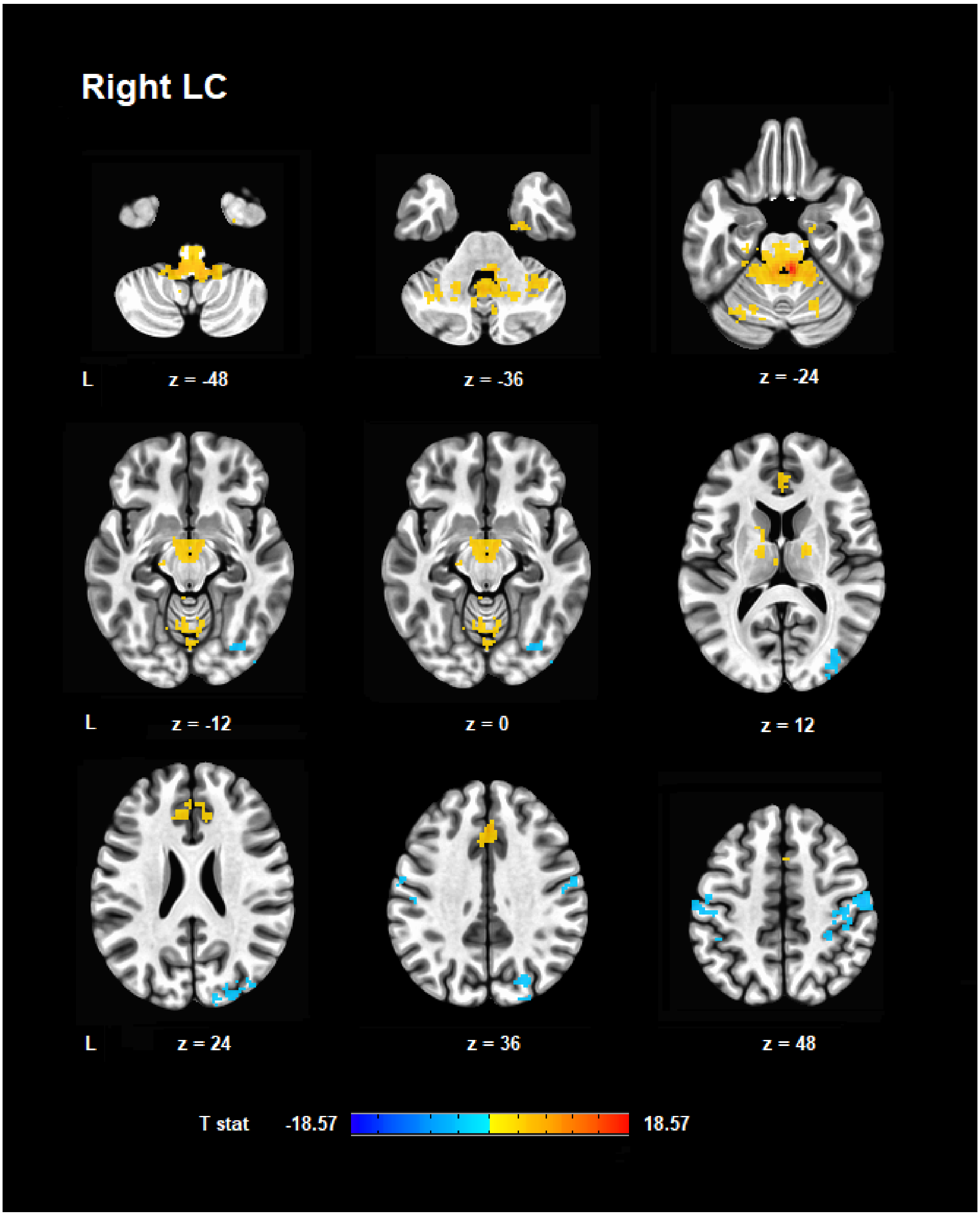


**Supplementary Figure S5.** The functional connectivity patterns of the right LC assessed with cluster-level FWE ($p < 0.05$) following cluster-forming threshold at uncorrected $p < 0.001$. Warm and cold colours indicate, respectively, positive and negative correlations.

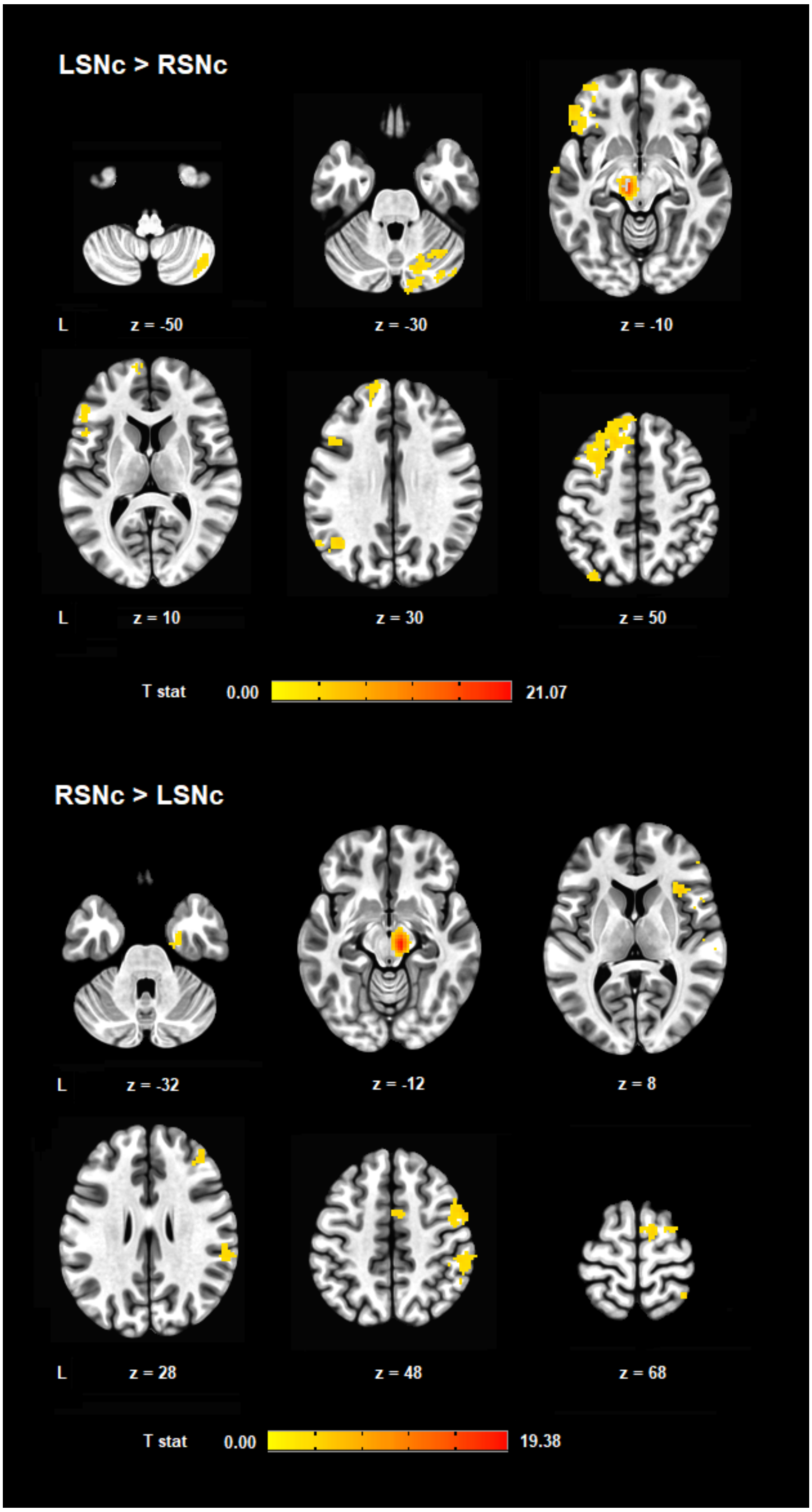


**Supplementary Figure S6.** Brain areas with differential functional connectivity with the left and right SNc. The results are corrected at the voxel-level with false discovery rate (FDR) < 0.05, and the cluster-forming threshold (k) is set at 40 voxels. Regions with more positive correlation with the left SNc are shown in the panel above, whereas clusters with more positive coupling to the right SNc are depicted in the panel below.

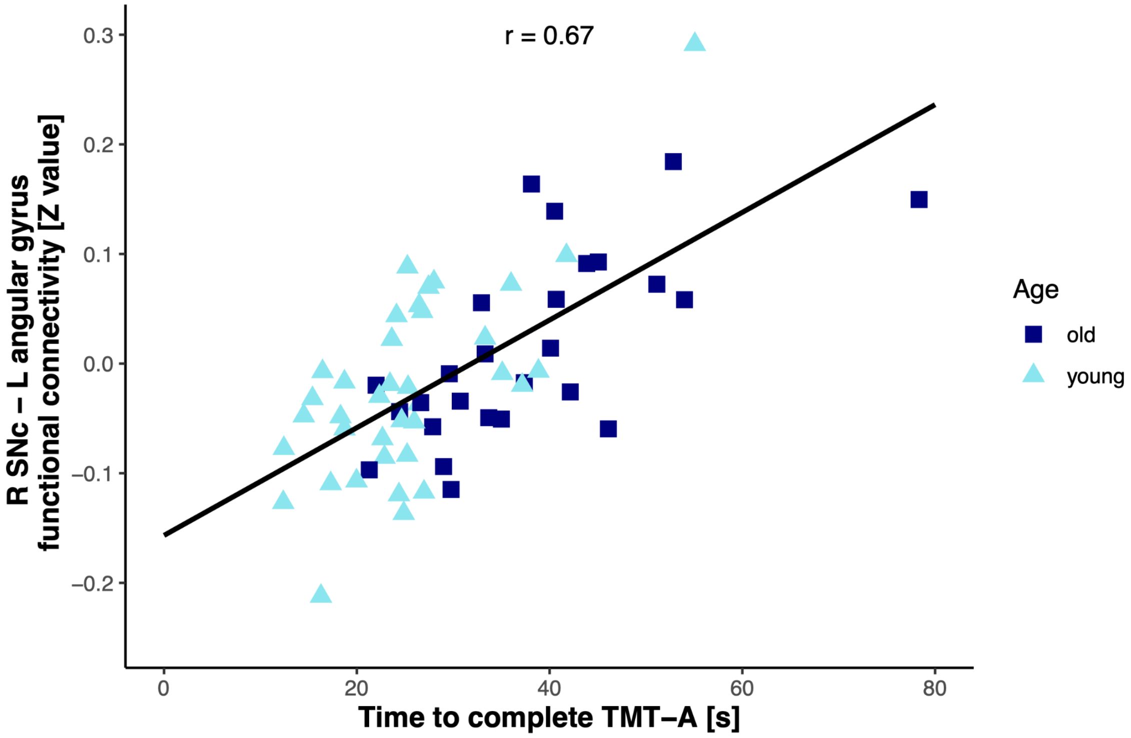


**Supplementary Figure S7.** Correlation plot of the TMT-A task completion time and resting-state functional connectivity between the right SNc and the left angular gyrus. Dark blue squares indicate participants assigned to the elderly group, while light blue triangles represent the younger group.

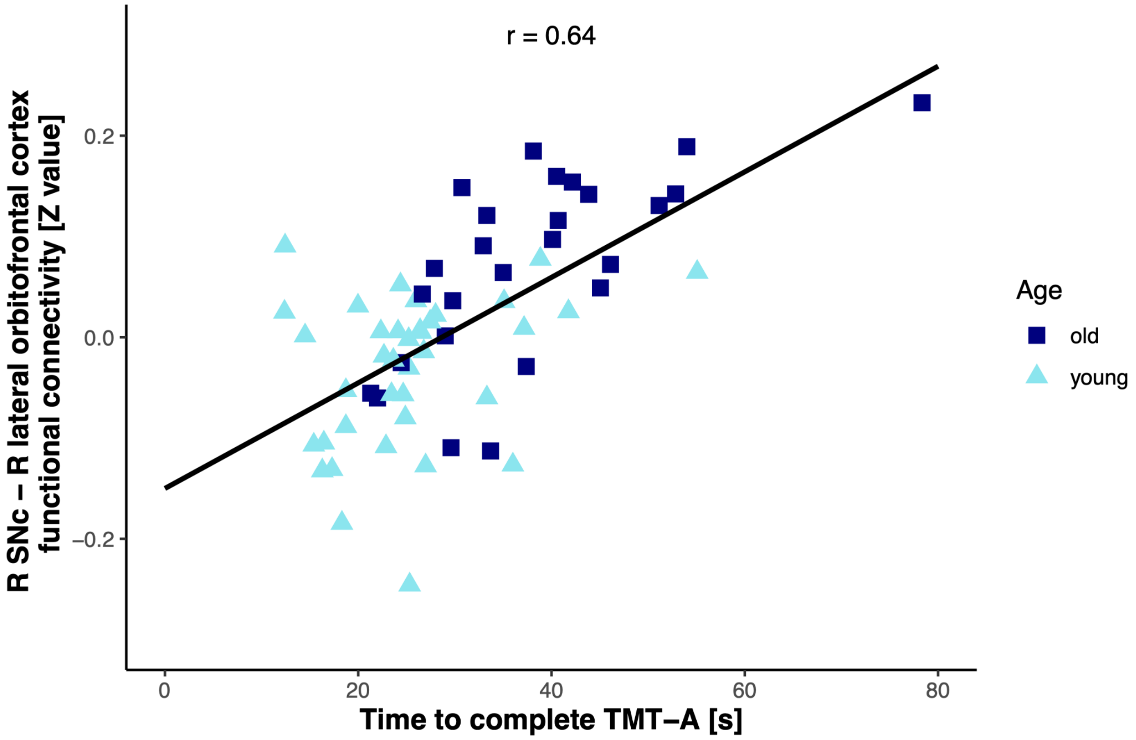


**Supplementary Figure S8.** Correlation plot of the TMT-A task completion time and resting-state functional connectivity between the right SNc and the right lateral orbitofrontal cortex. Dark blue squares indicate participants assigned to the elderly group, while light blue triangles represent the younger group.

**Supplementary Table S1.** The differences in whole-brain resting-state functional connectivity of the left and the right substantia nigra pars compacta (SNc). Paired t-test, voxel-level FDR < 0.05 and minimal cluster extent of 40 voxels.

| MNI | Location | Voxels | T-stat |
|---|---|---|---|
| *L SNc > R SNc* | | | |
| -18, 28, 50 | L superior frontal gyrus<br>L supplementary motor area<br>L dorsomedial prefrontal cortex<br>L middle frontal gyrus<br>L subgenual cingulate cortex | 1485 | 5.94 |
| -38, 42, 4 | L inferior frontal gyrus pars triangularis<br>L inferior frontal gyrus pars orbitalis<br>L lateral orbitofrontal cortex | 650 | 5.92 |
| 12, -84, -26 | R cerebellar lobule VIIa crus I | 304 | 5.34 |
| -9, -22, -12 | L midbrain | 286 | 21.07 |
| -41, -63, 29 | L angular gyrus | 157 | 5.12 |
| 44, -63, -51 | R cerebellar lobule VIIa crus II | 87 | 5.14 |
| -31, -77, 47 | L angular gyrus | 84 | 4.30 |
| 35, -63, -33 | R cerebellar lobule VIIa crus I | 51 | 5.40 |
| -61, -4, -15 | L middle temporal gyrus | 49 | 4.58 |
| -66, -28, -7 | L middle temporal gyrus | 47 | 4.10 |
| *R SNc > L SNc* | | | |
| 60, -27, 20 | R supramarginal gyrus<br>R precentral gyrus | 452 | 5.32 |
| 35, 22, 8 | R insula<br>R inferior frontal gyrus pars opercularis | 347 | 5.90 |
| 10, -20, -10 | R midbrain | 295 | 19.38 |
| 56, 3, 45 | R precentral gyrus | 146 | 5.66 |
| 56, -59, -3 | R inferior temporal gyrus<br>R middle temporal gyrus | 95 | 5.17 |
| 12, -1, 68 | R supplementary motor area | 91 | 4.61 |
| 5, 1, 47 | R supplementary motor area | 79 | 4.32 |
| 40, -52, 66 | R superior parietal lobe | 69 | 5.42 |
| 26, -8, -37 | R entorhinal cortex | 58 | 3.92 |

| | | | |
|---|---|---|---|
| 35, 47, 18 | R dorsolateral prefrontal cortex | 41 | 4.21 |
| 42, 40, 31 | R dorsolateral prefrontal cortex | 40 | 4.66 |

**Supplementary Table S2.** The comparison of the behavioural performance between the groups.

| Behavioural measure | Young | | Old | | F-stat | FDR |
|---|---|---|---|---|---|---|
| | Mean[a] Median[b] | SD[a] Range[b] | Mean[a] Median[b] | SD[a] Range[b] | | |
| TAP Alertness *mean reaction time [ms]* | 216[b] | 178 - 280.5[b] | 245[b] | 191 – 363.5[b] | 11.82 | 0.002 |
| TMT-A *completion time [s]* | 24.65[b] | 12.37 – 55.07[b] | 36.19[b] | 21.29 – 78.34[b] | 33.42 | < 0.001 |
| TMT-B - TMT-A *completion time [s]* | 24.43[b] | 3.22 – 79.88[b] | 42.46[b] | 6.54– 100[b] | 17.73 | < 0.001 |
| TAP Working Memory *percentage of correct responses [%]* | 93.33[b] | 14.29 – 100[b] | 80[b] | 10.71 – 100[b] | 6.50 | 0.020 |
| TAP Working Memory *mean correct response time [ms]* | 525[b] | 92 – 926[b] | 575[b] | 395 – 1244[b] | 3.46 | 0.081 |
| BAS Reward Responsiveness *total score [points]* | 17.09[a] | 2.04[a] | 16.50[a] | 2.18[a] | 1.07 | 0.306 |

[a] Means and standard deviations (SDs) indicate the use of parametric statistics.

[b] Medians and ranges indicate the use of non-parametric tests.
